# Curriculum Learning: sequential acquisition of task complexity enhances neuronal discriminability

**DOI:** 10.1101/2024.09.10.612027

**Authors:** Maria Shujah, Suyash Joshi, Mari Nakamura, Tania Rinaldi Barkat

## Abstract

Curriculum Learning (CL) is a strategy where concepts with increasing complexity are acquired sequentially. Despite its widespread application, its putative neural mechanisms are poorly understood. In this study, using a multi-staged Go/No-Go auditory discrimination paradigm for mice, we show that CL improves and accelerates learning in a unidirectional way. By combining behavioral data with *in vivo* electrophysiological recordings of the auditory cortex, we demonstrate that neuronal discriminability is enhanced with sequential training. This improvement suggests that CL involves a convergence of common discriminable features in the auditory stimuli shared across tasks of different complexity. Our findings uncover specific properties of CL at the level of neuronal discriminability, distinguishing it from other forms of learning strategies.

---

Throughout human history, the process of learning has been strategically designed to optimize efficiency. Sequential knowledge acquisition, where simpler tasks precede more complex ones, has proven to be an effective approach. For example, children are taught maths in a meaningful sequence of concepts where arithmetic precedes algebra and algebra precedes calculus^1,2^. This phenomenon, known as Curriculum Learning (CL), accelerates learning a complex task by abstracting the knowledge learned from easier tasks ^3,4^. Similarly, in machine learning, the approach of gradually training models on simple to complex datasets has demonstrated substantially improved performance and accelerated training process ^5-7^.

CL leverages the brain’s ability to form adaptable low-dimensional mental models or ‘schemas,’ which encode learning rules and features. These schemas allow new experiences to modify or refine them, enhancing subsequent learning ^8-13^. While rule learning has been extensively studied in rodents^14-17^, primates^18^, and humans^19,20^, the specific strategy of progressively applying learned rules from simpler tasks to more challenging ones has neither been clearly distinguished nor thoroughly investigated. Our study addresses this oversight by exploring both the behavioral strategy of CL and its neural counterparts, aiming to elucidate why this progressive approach significantly enhances learning efficiency.

To systematically characterize CL behavior, we designed a multi-staged appetitive Go/No-Go learning paradigm where a hard auditory discrimination task was preceded by an easier one (Fig. 1a, b, c). Both tasks required the mice to discriminate between a pulse train with a single tone (No-Go) and a pulse train with alternating tones (Go). The key difference between the tasks was the nature of the tones: the easy task used pure frequency tones (PT) (F^0^), while the hard task used harmonic complex tones (HCT) 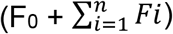 (Fig. 1d). Despite the increased complexity of the auditory stimuli, the fundamental structure—alternating vs. non-alternating F^0^s—remained consistent across both tasks.

**Fig. 1.**
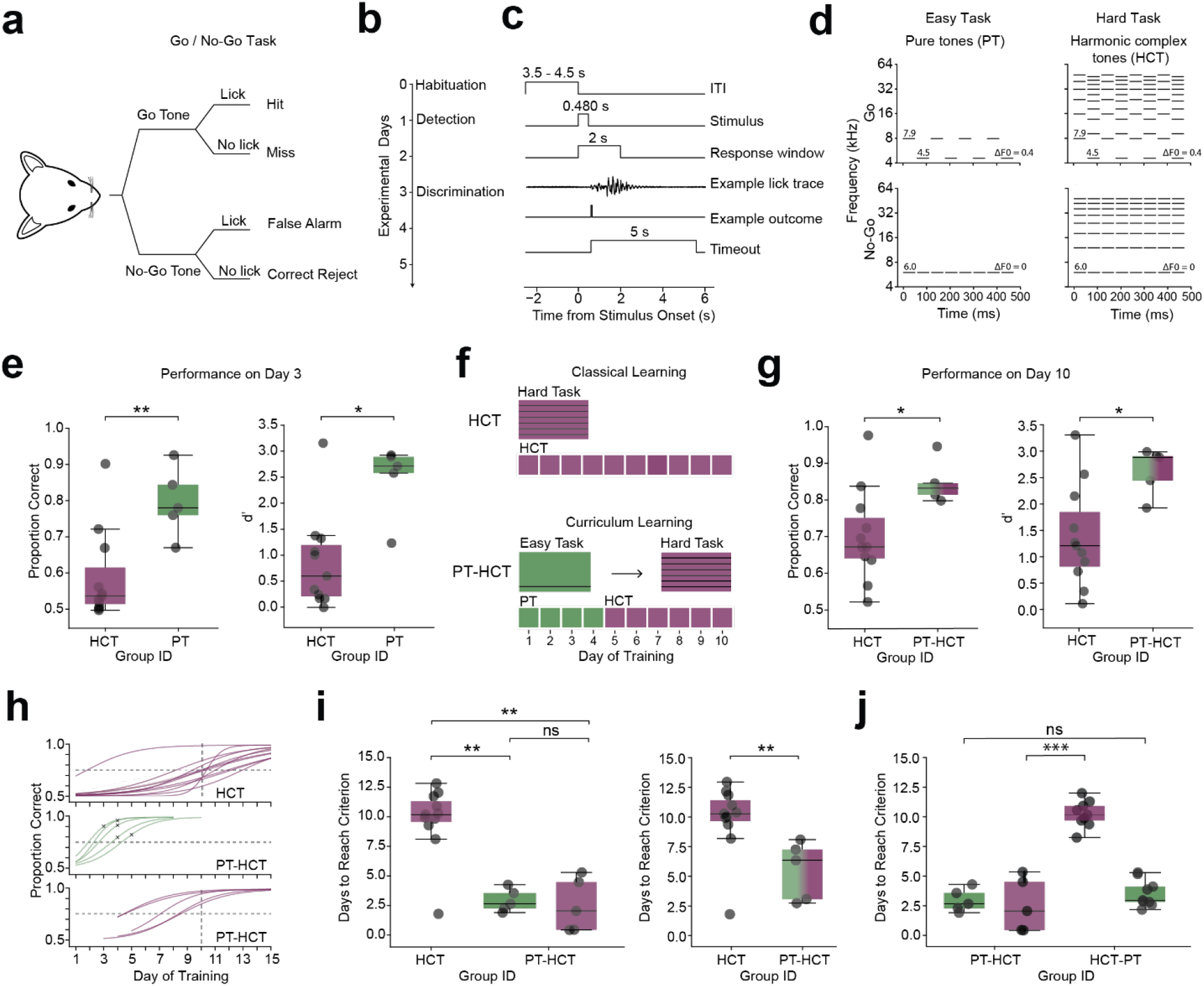
Curriculum learning enhances and accelerates auditory discrimination learning. **a**, Schematic of the Go/No-Go auditory discrimination task. **b**, Sequential training phases starting with Habituation, followed by Tone Detection (lick response to tone), and culminating in Tone Discrimination (discrimination between Go and No-Go tones). **c**, Sequence of events in a trial, starting with a random inter-trial interval (ITI), a 480 ms stimulus presentation, followed by a 2 s response window where a lick triggers an outcome which is either a reward (sweetened soymilk drop) or a penalty (air puff and 5 s timeout). **d**, Schematic for sounds used in the tasks: Easy task: Go, eight pulses of two alternating pure tones; No-Go, eight pulses of one pure tone. Hard task: Go, eight pulses of alternating harmonic complex tones with two fundamental frequencies (F_0_); No-Go, eight pulses of non-alternating harmonic complex tones with a single F_0_. **e**, Performance metrics on day 3 for the hard (HCT, purple) tasks and the easy (PT, green), showing percentage correct and d-prime (d’) values. n = 11 mice (Group HCT), n = 5 mice (Group PT), p=0.0087 for percent correct, p=0.019 for d-prime, Mann-Whitney test. **f**, Training regimens for two cohorts: Top: Group HCT, trained only on the Hard task; Bottom: Group PT-HCT, initially trained on the Easy task until achieving 75% accuracy, then advancing to the Hard task. **g**, Comparative performance of Group HCT (purple) and Group PT-HCT (green-purple gradient) on day 10 of training in terms of percentage correct (left) and d-prime (right). Number of mice as in (e), p = 0.027 for percent correct, p = 0.027 for d-prime, Mann-Whitney test. **h**, Learning kinetics across tasks for Group HCT (top) and Group PT-HCT (middle and bottom). The horizontal grey line denotes 75% accuracy (performance criterion); the vertical grey line indicates Day 10. Each curve represents a mouse. The indicators (x) in the middle plot indicate the day of training at which the mice were moved to HCT training. For the top and bottom plots, the curves beyond Day 10 are extrapolated based on the data up to Day 10, as training concluded on this day. For the middle plot, the curves beyond the days indicated with x are extrapolated as the mice progressed to the next task after this day. **i, (**Left) Days required to reach the 75% performance criterion in individual tasks by Group HCT (purple) and PT-HCT (green-purple gradient). Number of mice as in (e), p=0.0087 for Group HCT task HCT vs. Group PT-HCT task PT, 0.84 for Group PT-HCT task PT vs. Group PT-HCT task HCT, 0.0032 for Group HCT task HCT vs. Group PT-HCT task HCT, Mann-Whitney test. (Right) Days required to reach the 75% performance criterion in the entire training by Group HCT (purple) and PT-HCT (green-purple gradient). Number of mice as in (e), p=0.0087, Mann-Whitney test. **j**, Days to reach criterion in individual tasks by Group PT-HCT and Group HCT-PT. n = 5 (Group PT), n = 9 (Group HCT-PT), p = 0.0009 for Group PT-HCT task HCT vs. Group HCT-PT task HCT, p = 0.36 for Group PT-HCT task PT vs. Group HCT-PT task PT, Mann-Whitney test. Data are based on psychometric functions fitted with a logistic model. In the boxplots, lines represent the median, 25th (first quartile, Q1), and 75th percentiles (third quartile, Q3), whiskers denote the highest and lowest values within 1.5 times the interquartile range (IQR) from Q1 and Q3.

To test the perceptual difficulty of the two tasks, we divided the mice into two groups: one group was trained exclusively on PT discrimination, while the other was trained on HCT discrimination. By the third day of training, a marked difference in performance was observed: the PT group exhibited significantly higher performance than the HCT group, demonstrating that PT discrimination is easier than HCT discrimination (Fig. 1e).

To study the effect of CL, we then created two cohorts: one cohort underwent progressive training from the easy task (PT discrimination) to the hard task (HCT discrimination) after having achieved a predefined accuracy criterion of 75% on the easy task (PT-HCT group). In contrast, the second cohort was subjected to the hard task from the outset (HCT group) (Fig. 1f). Assessment on the tenth day of training revealed that the PT-HCT group outperformed the HCT group significantly (Fig. 1g). Moreover, it also reached the performance criterion in fewer days (Fig. 1h, i).

To test if the performance improvements were task-sequence dependent, we trained a third group first on the hard task and later on the easy task (HCT-PT). Surprisingly, training on the hard task first did not significantly improve performance on the easy task compared to no prior training (Fig. 1j). This finding highlights the unidirectional effect of CL, distinguishing it from the broader concept of generalization ^21^.

To explore the neuronal underpinnings of CL, we conducted acute *in vivo* electrophysiological recordings from the primary auditory cortex (A1). A1 is crucial for processing complex sounds and higher cognitive functions, particularly learning, making it ideal for investigating the neural mechanisms of CL ^22-26^. We recruited three mouse cohorts: naïve, trained on PT discrimination only (group PT), and trained sequentially on PT followed by HCT discrimination (group PT-HCT) (Fig. 2a, b). The recordings were done at the last day of training of each training group in awake, passive-listening mice. We hypothesized that neuronal responses would reflect the difficulty levels of the easy and the hard task (Fig. 2c). If PT discrimination is easier than HCT discrimination, it is pertinent to expect that the neuronal responses to PTs (Go and No-Go) are less similar than to HCTs (Go and No-Go).

**Fig. 2.**
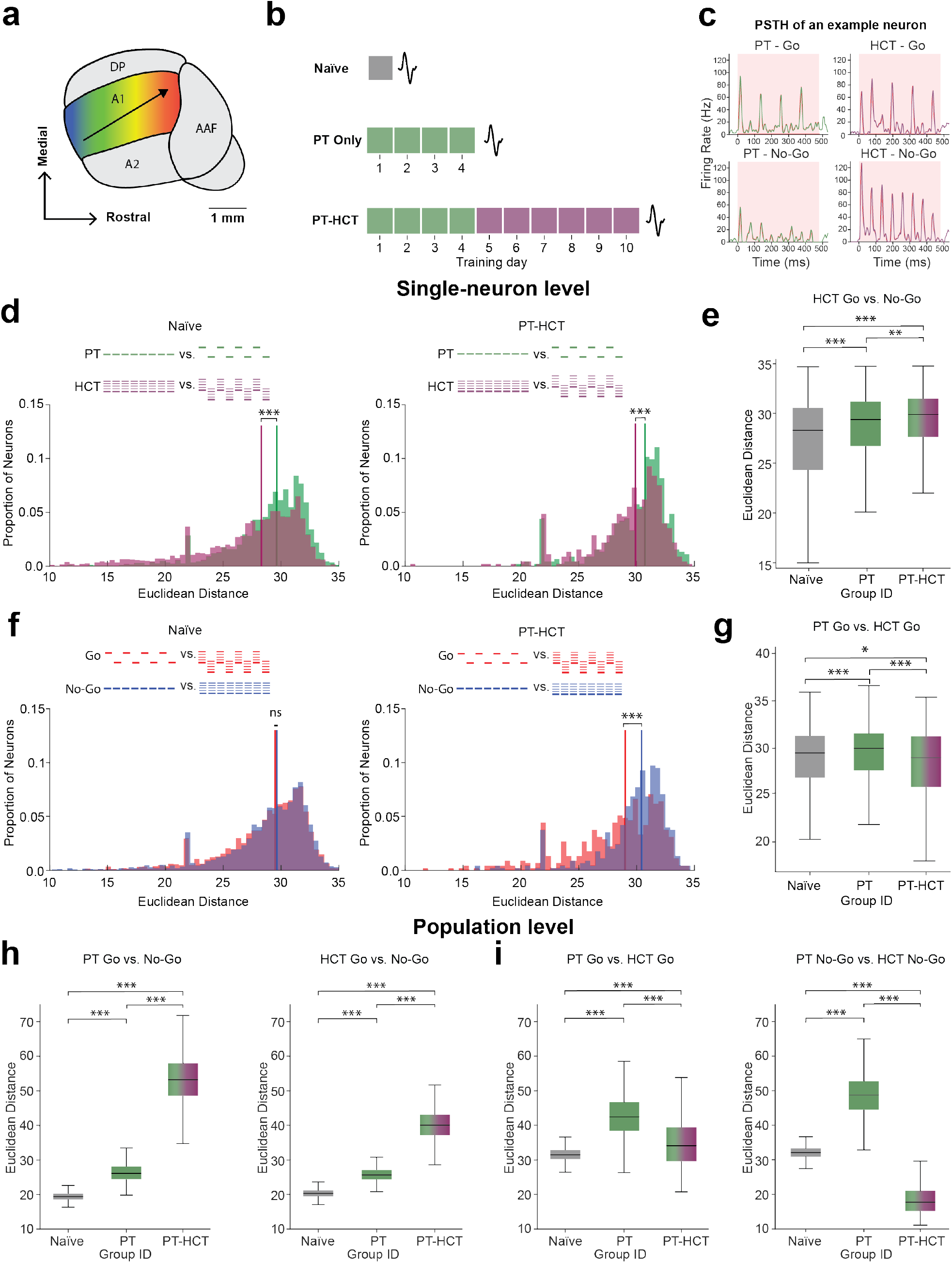
Training on the easy task alone enhances neuronal discriminability for tones used in easy and hard tasks. **a**, Schematic of the auditory cortex, highlighting the tonotopic organization within the primary auditory cortex (A1) and adjacent regions: secondary auditory cortex (A2), Anterior Auditory Field (AAF), and Dorsoposterior (DP). **b**, Overview of training groups: Naïve, PT and PT-HCT. Electrophysiological recordings were conducted acutely on the final training day as mice passively listened to the auditory stimuli from both PT and HCT discrimination tasks. **c**, Example Peri-Stimulus Time Histograms (PSTH) in response to Go and No-Go sounds of both tasks in a single neuron. The shaded area indicates stimulus duration (480 ms). **d**, (Top) Auditory stimuli used for response comparison. Histograms depict the Euclidean distances between single-neuron responses to PT Go vs. PT No-Go (green) and HCT Go vs. HCT No-Go (purple) across groups: Naïve (left) and PT-HCT (right). n=2664 units (Naïve), n= 733 units (PT), n=453 units (PT-HCT), p < 0.0001 (Naïve), p = 0.0002 (PT-HCT), Mann Whitney Test. **e**, Single neuron-level Euclidean distances for HCT Go vs. HCT No-Go for all three training groups. Number of units as in (d), p = 0.00035(Naïve vs. PT), p < 0.0001(Naïve vs. PT-HCT), and p = 0.02 (PT vs. PT-HCT), Kruskal-Wallis test followed by Dunn’s post-hoc test. **f**, (Top) Auditory stimuli used for response comparison. Histograms show single-neuron Euclidean distances for PT Go vs. HCT Go (red) and PT No-Go vs. HCT No-Go (blue) across groups: : Naïve (left) and PT-HCT (right). Number of units as in (d), p = 0.059 (Naïve), p < 0.0001 (PT-HCT), Mann Whitney Test. The median Euclidean distance is marked by the straight line in both histograms. **g**, Single neuron-level Euclidean distances for PT Go vs. HCT Go in all three training groups. p < 0.0001(Naïve vs. PT and PT vs. PT-HCT). Number of units as in (d), p = 0.04 (Naïve vs. PT-HCT), Kruskal-Wallis test, followed by Dunn’s post-hoc test. **h**, Population-level Euclidean distances for PT Go vs. PT No-Go (left) and HCT Go vs. HCT No-Go (right). Population responses are based on bootstrapped resampling of single neuron responses (x1000). n = 1000 for each group (Naïve, PT, PT-HCT), p < 0.0001 for all comparisons, Permutation test. **i**, Population-level Euclidean distances between PT Go vs. HCT Go (left) and PT No-Go vs. HCT No-Go (right), also derived from bootstrapped data. Number of samples as in (h), p < 0.0001 for all comparisons, Permutation test. In the histograms, lines represent the median. In the boxplots, lines represent the median, 25th (first quartile, Q1), and 75th percentiles (third quartile, Q3), whiskers denote the highest and lowest values within 1.5 times the interquartile range (IQR) from Q1 and Q3.

We assessed neuronal discriminability by measuring the Euclidean distance between neuronal responses to Go and No-Go sounds for both task types. This metric reflects the similarity between neuronal firing patterns—shorter distances indicate higher similarity and thus poorer discriminability. At the single-neuron level, we found that in the naïve group, the Euclidean distance between responses to PTs was significantly larger than those for HCTs, aligning with the behavioral observation that PTs are easier to discriminate than HCTs. As might have been expected, after PT-HCT training, the Euclidean distance between HCTs increased significantly, indicating enhanced learned discriminability (Fig. 2d). Notably, in the PT group, discriminability for HCTs (not trained on) was also significantly improved compared to the naïve group, suggesting that training on the PTs alone enhances neuronal discriminability for the HCTs (Fig. 2e).

We hypothesized that the enhancement in discriminability could be attributed to the common schemas used between the Go and/or No-Go sounds of the two tasks, leading to more similar neuronal responses. If this hypothesis is correct, we would expect an increased similarity in neuronal responses to both Go and No-Go sounds across tasks. To explore this, we examined the Euclidean distance between Go sounds (PT Go vs. HCT Go) and No-Go sounds (PT No-Go vs. HCT No-Go) across all three groups. Our findings indicate that, specifically in the PT-HCT group, the responses to the Go sounds became more similar following training (Fig. 2f, g). Extending our analysis to population-level activity to assess consistency with single-neuron level activity, we found that the similarity patterns generally aligned with single-neuron observations. However, at the population level in the PT-HCT group, it was the No-Go sounds that exhibited increased similarity across tasks, contrasting with the single-neuron level where the Go sounds showed greater similarity (Fig. 2h, i).

To decipher which sound features were abstracted across tasks, we analyzed the raw population-level Peri-Stimulus Time Histograms (PSTH) for each sound. In both naïve and PT groups, the PSTH prominently reflected the pulse structure of the sounds, with peaks corresponding to each pulse onset. In contrast, the PT-HCT group’s PSTH structure shifted, weakly reflecting each pulse onset while more distinctly representing a binary classification between alternating Go (higher firing rate) and non-alternating No-Go (lower firing rate) sounds (Fig. 3a). This shift led us to hypothesize that with training, neurons begin to abstract features relevant for discrimination that are shared between the two tasks, e.g. alternation or F^0^.

**Fig. 3.**
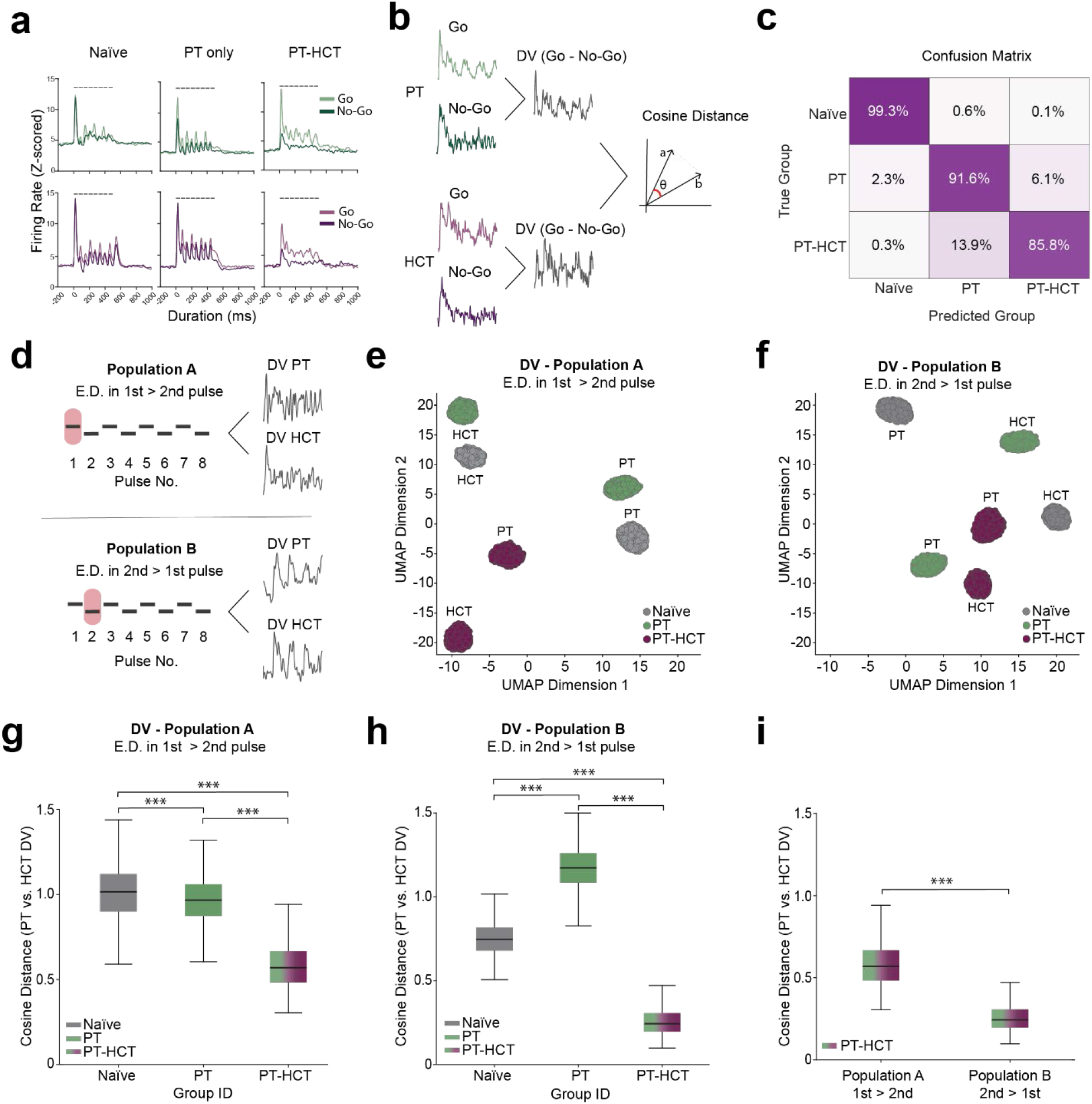
Curriculum learning leads to feature abstraction across easy and hard tasks. **a**, Example population PSTH shows a change in pattern across training groups. A schematic of the pulse train is displayed on top of each plot. **b**, Procedure for calculating discriminability vectors: The difference between PT Go vs. No-Go and HCT Go vs. No-Go population responses provide discriminability vectors ‘a’ and ‘b’ respectively. Population responses are based on bootstrapped resampling of single neuron responses (x1000). Cosine distance is used to quantify the similarity between these vectors, with lower values indicating higher similarity. DVs are calculated using bootstrapped population activity data. Traces are PSTHs from example neurons. **c**, Confusion matrix from an SVM classifier trained to predict the training group using discriminability vectors. True groups are on the Y-axis; predicted groups are on the X-axis. Each cell shows the percentage of total predictions where the predicted group (column) aligns with the true group (row). n = 1000. **d**, Strategy to identify two neuronal populations. Population A: Neurons with a higher Euclidean distance between Go and No-Go during the first pulse compared to the second pulse. Population B: Neurons with a higher Euclidean distance during the second pulse (after 60 ms from stimulus onset) compared to the first. Traces are DVs from example neurons. **e, f**, UMAP visualization of DVs from Population A and B, respectively, distinguishing the three training groups: Naïve, PT, and PT-HCT, shown in grey, green, and purple, respectively. **g, h**, Cosine distances between DVs from Population A and B, respectively, for all three training groups. n = 1000 for each group (Naïve, PT, PT-HCT), p < 0.0001 for all comparisons, Permutation tests. **i**, Cosine distances between DVs from Population A and B, respectively, for Group PT-HCT, showing significantly lower Cosine distance between DVs from Population B compared to Population A. n = 1000 for each group (Naïve, PT, PT-HCT), p < 0.0001 for all comparisons, Permutation tests. In the boxplots, lines represent the median, 25th (first quartile, Q1), and 75th percentiles (third quartile, Q3), whiskers denote the highest and lowest values within 1.5 times the interquartile range (IQR) from Q1 and Q3.

To test our hypothesis, we first computed discriminability vectors (DV, difference between Go vs. No-Go PSTH) for neuronal population activity to see if distinct, characteristic features emerged within the neuronal activity patterns of each training group as seen in the raw PSTHs (Fig. 3b). We employed a Support Vector Machine (SVM) to analyze the decoding accuracy of the DVs derived from each training group. This machine learning model was trained to classify the vectors, aiming to decode which training group (Naïve, PT, or PT-HCT) a given DV belonged to. The decoding accuracy was remarkably high (Fig. 3c).

Having established the distinctness of the DVs, we next investigated which feature is abstracted across the two tasks. We distinguished two distinct neuronal populations (see Methods). Population A consisted of neurons that showed enhanced discriminability (greater Euclidean distance between Go and No-Go) during the first pulse compared to the second for both PT and HCT discrimination. This is likely due to F_0_ discrimination. Population A proportion of total neurons for each group are as follows: naïve (0.22), PT (0.19), PT-HCT (0.20). Population B had higher discriminability during the second pulse, coinciding with the first alternation in the Go sounds, for both PT and HCT discrimination. This could be due to alternation pattern discrimination. Population B proportion of total neurons for each group are as follows: naïve (0.29), PT (0.25), PT-HCT (0.29) (Fig. 3d). Plotting the discriminability vectors (DVs) in a lower-dimensional space using UMAP revealed that the DVs from both populations cluster more closely for the PT-HCT group (Fig. 3e, f). We then evaluated the directional similarity between the DVs for both tasks using the Cosine distance metric, where shorter distance indicates higher similarity. Notably, for both populations, the DVs from both tasks exhibited significantly greater similarity within the PT-HCT group (Fig. 3g-h), suggesting that training induces the utilization of shared discriminative features. However, Population B exhibited greater similarity than Population A indicating that the degree of abstraction increased more after the second pulse (Fig. 3i). While the precise feature used for discrimination across the tasks cannot be definitively determined, this result suggests that the alternation pattern is a key feature abstracted through CL.

In this study, we aimed at identifying what neuronal mechanisms support the effect of performance improvement due to CL. Our results demonstrate that sequentially structuring auditory learning tasks with increasing complexity significantly enhances both the speed and accuracy of learning. Importantly, this enhancement is unidirectional. *In vivo* electrophysiology revealed that neuronal discriminability improves with training and that training on the easy task alone enhances discriminability for the hard task, at both single-cell and population levels. The differences between Go and No-Go responses were distinct across different training groups. However, within the PT-HCT group, these differences, for tones used both in the easy and hard tasks, became highly similar, suggesting a convergence on common discriminative features.

The findings of this study are significant for several reasons: Firstly, they provide a clear behavioral paradigm for studying CL, distinguishing it from classical learning strategies that lack a sequential framework, as well as from generalization, which involves bidirectional rather than unidirectional transfer of learning. Secondly, our electrophysiological data reveal the distinct neuronal changes that occur when training in a sequential manner. Specifically, we observe high-level abstraction of the relevant feature(s) across the two tasks, suggesting the formation of cognitive schemas. This identifies the putative neural mechanisms that enhance learning efficiency in CL.

While further research is needed to understand the precise computational processes involved, our work lays the groundwork for future studies on the dynamic evolution of feature abstraction in progressively challenging tasks. It also invites exploration into how the structure and content of curricula can differentially impact learning efficiency, consistent with recent findings on how CL can uncover learning principles ^27^. These insights underscore the benefits of structuring educational and training programs to align with the inherent learning processes of the brain.

## Methods

### Animals

All animal procedures were performed in accordance with the University of Basel animal care and use guidelines and were approved by the Veterinary Office of the Canton Basel-Stadt, Switzerland. C57BL/6JRj mice (Janvier, France) were used for the experiments. Both male and female mice, a minimum of 42 days (6 weeks) of age were used for this study. Mice were housed in a 12-hour light/dark cycle with free access to food and water. All mice were single-housed and food-restricted during the behavior training period to maintain 10-15% body weight loss. Food was provided in calculated amounts after every training session. Experiments were performed in the light phase of the daily cycle.

### Surgical procedure

Mice were anesthetized with isoflurane in O_2_ (4% induction, 1.2 to 2.5% maintenance), and local analgesia was provided with subcutaneous injection of bupivacaine/lidocaine (0.01 mg/animal and 0.04 mg/animal, respectively). The depth of anesthesia was monitored by the breathing rate and absence of pinch withdrawal reflex. Body temperature was maintained at 37 °C via a heating pad (FHC, ME, USA), and lubricant ophthalmic ointment was applied on both eyes. A custom-made stainless-steel head-restraint post was fixed on the bone on top of the left hemisphere and used to head-fix the animals. Using a scalpel, a craniotomy (∼2×2 mm^2^) was performed just above the auditory cortex. The dura was left intact and was covered with silicone oil to prevent drying. The craniotomy was covered with a thick layer of Kwik-Cast Sealant (WPI, Sarasota, FL, USA) to protect it from external agents. Mice were placed back in their home cage for 2 hours to recover from anesthesia.

### Behavioral task

Mice were trained to lick in response to a sound stimulus, within a response window of 2 seconds from sound onset. On licking in response to a Go tone, mice received a drop of sweetened, vanilla-flavored soy milk (Alpro) and the response was counted as a ‘Hit’. If the mouse did not respond to the Go tone, the trial was counted as a ‘Miss’. On licking in response to a No-Go tone, mice received a punishment of a brief air puff delivered near the eye followed by a time-out of 5s, and the trial was considered a ‘False Alarm’. If the mouse did not lick in response to the No-Go tone, the trial was counted as a ‘Correct Reject’. Sounds were delivered without preceding cues at random interstimulus intervals (ITI) ranging from 3.5 to 4.5 s. Licks were detected with a piezo sensor attached to the reward spout.

### Habituation

On the first day of training, mice were habituated with the experimental rig, and were presented with 25 trials. These stimuli consisted of a pulse train of white noise, followed by an automatic dispensing of reward.

### Tone Detection

Next, mice were trained to lick in response to a sound. No punishment was given at this stage. Mice moved to the next stage when the Hit rate was above 90%.

### Tone Discrimination

At this stage, mice were trained to discriminate between a Go and a No-Go tone. Each session consisted of 500 trials in pseudorandom order (250 Go trials, 250 No-Go trials). For each trial, the probability of a Go and No-Go stimulus presentation was 50%. Stimulus, reward delivery, and behavioral data acquisition were controlled by a personal computer through RPvdsEx real-time design studio and an RZ6 multifunction processor (Tucker-Davis Technologies).

### Auditory stimulation

All auditory stimuli were composed of a train of eight pulses, each 50 ms in duration, with 4 ms onset and offset ramps shaped by a cosine squared function. Pulses were interspersed with 10 ms of silence. Sounds were delivered in a free-field setting using speakers located 10 cm from the head-fixed animal’s ears, at a consistent sound pressure level of 60 dB SPL.

In the Easy Task, the PT Go were eight pulses alternating between F_0_s of 7.917 kHz and 4.547 kHz, representing a difference of 0.4 octaves around 6 kHz while the PT No-Go consisted of eight identical pulses at an F_0_ of 6 kHz. In the Hard Task, the HCT Go alternated between HCT composed of F_0_s of 4.547 kHz and 7.917 kHz within the 4 kHz to 50 kHz range, all in phase. The HCT No-Go stimuli were eight pulses of an HCT summing all harmonics of an F_0_ of 6 kHz within the 4 kHz to 50 kHz range, all in phase. We acknowledge that the term “fundamental frequency” is technically applicable only to HCTs and not to PTs. However, for the sake of simplicity and better comprehension, we use the term “fundamental frequency” consistently for both PTs and HCTs throughout this manuscript.

### *In vivo* electrophysiology

After a 20–24-hour recovery period from surgery, mice were head-fixed. The craniotomy was exposed and covered with silicon oil. A 4×16 electrode (A4 × 16-5 mm-50-200-177-A64, Neuronexus, MI, USA) was inserted into A1 with a motorized stereotaxic micromanipulator (DMA-1511, Narishige, Japan) at a depth of 952 ± 37 μm from pia, traversing the superficial, input, and deep layers. Recordings from A1 were confirmed in each penetration by increasing averaged BF from the most caudal to the most rostral shaft of the 4-shaft electrodes, confirming the tonotopic organization typical of A1. All recordings were performed in a sound-attenuating chamber (modified MAC-2 chambers, Industrial Acoustics Company Nordics, Hvidorve, Denmark). At the end of the recording sessions, each mouse was sacrificed with an intraperitoneal (i.p.) injection of pentobarbital followed by cervical dislocation.

## Data analysis

### Behavioral performance analysis

To assess the performance of mice in the auditory discrimination tasks, we calculated *d*′ (d-prime), a metric from signal detection theory that quantifies sensitivity to distinguish between signal (hits) and noise (false alarms). *d*′ is defined as:

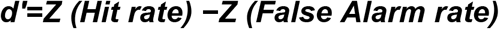

where *Z* is the inverse cumulative distribution function of the normal distribution. The Hit rate is the proportion of correct responses to Go stimuli, and the False Alarm rate is the proportion of incorrect responses to No-Go stimuli. Values of Hit and False Alarm rates were truncated between 0.0001 and 0.9999.

To calculate the number of days required for mice to reach the performance criterion, we fitted constrained psychometric functions to the behavioral data using the **psignifit** toolbox in MATLAB Mathworks, MA, USA). Specifically, we organized the data for each stimulus type (PT and HCT), with the x-axis representing the number of training days. The psychometric functions were fitted using a logistic model, with constraints applied to ensure the maximum performance did not exceed 95% correct responses and with a fixed lapse rate of 0.01. We generated predicted performance curves based on these fits and used interpolation to determine the number of days required for each mouse to achieve the performance criterion of 75% correct responses.

### In vivo electrophysiology Analysis

Extracellular recordings were digitized with a 64-channel recording system (RZ2 Bioamp processor, Tucker Davis Technologies, FL, USA) at 24414 Hz. Putative single and multi-units were identified offline using KiloSort version 2.5 (CortexLab, UCL, London, England; ^28^) followed by manual inspection of spike shape and signal-to-noise ratio, and auto- and cross-correlograms using Phy (CortexLab, UCL, London, England). Both single and multi-units were retained for analysis. Further analysis was performed using custom software in MATLAB.

### Metric space analysis

In our analysis, we quantitatively assessed the similarity between neural response patterns by calculating Euclidean and Cosine distances. These metrics were employed to measure the divergence and angular similarity between different stimuli responses. The calculations were performed using MATLAB.

Euclidean distance between two vectors, x and y, each representing neural activity (PSTH) during stimulus presentation in response to different training conditions, was computed as follows:

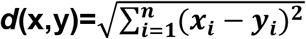

where *xi* and *yi* are the components of vectors x and y, respectively, and n is the number of components in each vector. This metric provides a measure of the “straight-line” distance between two points in an n-dimensional space.

Cosine distance measures the cosine of the angle between two vectors. It is derived from the cosine similarity, which is mathematically represented as:

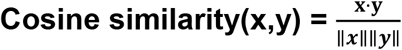

where ⋅ denotes the dot product of the vectors, and ∥**x**∥ and ∥**y**∥ are the normalized vectors. The Cosine distance is then calculated as:

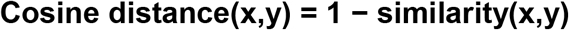

This metric ranges from 0 to 2, where 0 indicates no angular difference (high similarity) and 2 indicates an angle of 180 degrees between the vectors (maximum dissimilarity). It measures the orientation rather than the magnitude of the vectors, making it particularly useful for comparing normalized responses. These distances were computed using MATLAB’s built-in functions **pdist2** for Euclidean distance and **pdist** with the ‘cosine’ option for Cosine distance, ensuring efficient and standardized calculations.

For single neuronal response analysis, Euclidean distances were calculated between stimuli responses to PT Go vs. PT No-Go and HCT Go vs. No-Go for all training groups. Similarly, Euclidean distances between PT Go vs. HCT Go and PT No-Go vs. HCT No-Go were also calculated.

For the population response analysis, population responses were generated by bootstrap resampling (x 1000) of single-cell responses. Using these generated vectors, we calculated the Euclidean and distances between all stimuli comparisons.

### Discriminability Vector (DV) calculation and classification

Discriminability vectors were computed for both tasks. These vectors were calculated by taking the difference between the bootstrapped population responses to Go and No-Go stimuli.

The discriminability vector, **DV**, for each task is calculated using the equation:

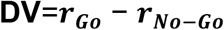

where:

1. *rGo* is the average population response vector to the Go stimuli.
2. *rNo—Go* is the average population response vector to the No-Go stimuli.

We calculated DVs for two neuronal populations, A and B. Population A consisted of neurons that have a higher Euclidean distance between both PT Go vs. No-Go and HCT Go vs. No-Go during the first pulse (0-60 ms) of the stimuli compared to the second (60-120 ms). The proportion of total neurons is as follows: Naïve (0.22), PT (0.19), PT-HCT (0.20). Population B consisted of neurons that had a higher Euclidean distance between PT Go vs. No-Go and HCT Go vs. No-Go during the second pulse (60 – 120 ms) of the stimuli compared to the first (0-60 ms). The proportion of total neurons is as follows: Naïve (0.29), PT (0.25), PT-HCT (0.29). For both populations, only neurons that met the Euclidean distance criteria for both PT and HCT tasks were selected.

To investigate the neural discriminability patterns across different training groups—Naïve, PT only, and PT-HCT—we employed a multiclass classification approach using a support vector machine (SVM) with error-correcting output codes (ECOC). This analysis was designed to determine how effectively the neural activity patterns reflected in the DVs could be used to predict the training group they belong to.

Prior to classification, the combined dataset was randomized to prevent order effects, ensuring an unbiased distribution of data points in training and testing datasets. We partitioned the data into training (70%) and testing (30%) sets using stratified sampling to maintain a proportional representation of each group. The SVM classifier was then trained on the training set using a linear kernel and a one-vs-one coding scheme for the ECOC model. Hyperparameter optimization was conducted via a 10-fold cross-validation within the training set, optimizing for model parameters such as box constraints and kernel scale. The optimization process aimed to minimize classification error and enhance generalization across unseen data.

Model performance was assessed on the held-out test set, calculating overall accuracy and group-specific precision and recall. A confusion matrix was generated to visualize the classifier’s performance across the three groups (Fig. 3c). The similarity between DVs was measured by calculating the Cosine distance between them (Fig 3g-i).

## Statistical tests

Statistics were conducted using MATLAB 2023b and Python 3. In the boxplots, lines represent the median, 25th (first quartile, Q1), and 75th percentiles (third quartile, Q3), while whiskers denote the highest and lowest values within 1.5 times the interquartile range (IQR) from Q1 and Q3, respectively. The Mann-Whitney U test was used to compare performance and days to reach the criterion between two training groups (Fig. 1d, f, g, i). For similarity analysis at the single-neuron level, the Mann-Whitney U test was used to calculate significance, with histogram lines representing the median value (Fig. 2d, f). For boxplots in this analysis, the Kruskal-Wallis test was employed followed by a post hoc Dunn-Sidak test (Fig. 2e, g). Given that the data for population-level similarity analysis were derived from bootstrapped resampling, a permutation test with 1000 permutations was used to test for significance between training groups. For DV similarity analysis, joint permutation tests were similarly used due to prior bootstrapping (Fig. 3g, h, i). All statistical tests used were two-sided. The effects were named significant if the p-value was smaller than 0.05 (*), 0.01 (**), and 0.001 (***) for a confidence interval of 95%, 99%, or 99.9%, respectively.

In this study, effect size and eta-squared were calculated to quantify the magnitude of observed effects. For the Mann-Whitney U Test, we calculated the rank-based effect size **r** as follows:

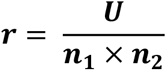

where U is the Mann-Whitney U statistic, representing the difference in ranks between two groups, and n1 and n2 are the sample sizes for Group 1 and Group 2, respectively. The value of **r** ranges between 0 and 1, where higher values indicate a larger effect size. For the Kruskal-Wallis Test, we calculated eta-squared (**η^2^**) to measure the proportion of variance explained by the group differences. Eta-squared is calculated as:

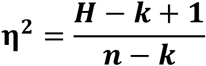

where H is the Kruskal-Wallis statistic, k is the number of groups and n is the total number of observations.

We did not calculate effect sizes for permutation tests on data that was bootstrapped due to the resampling procedures involved.

**Table 1.**
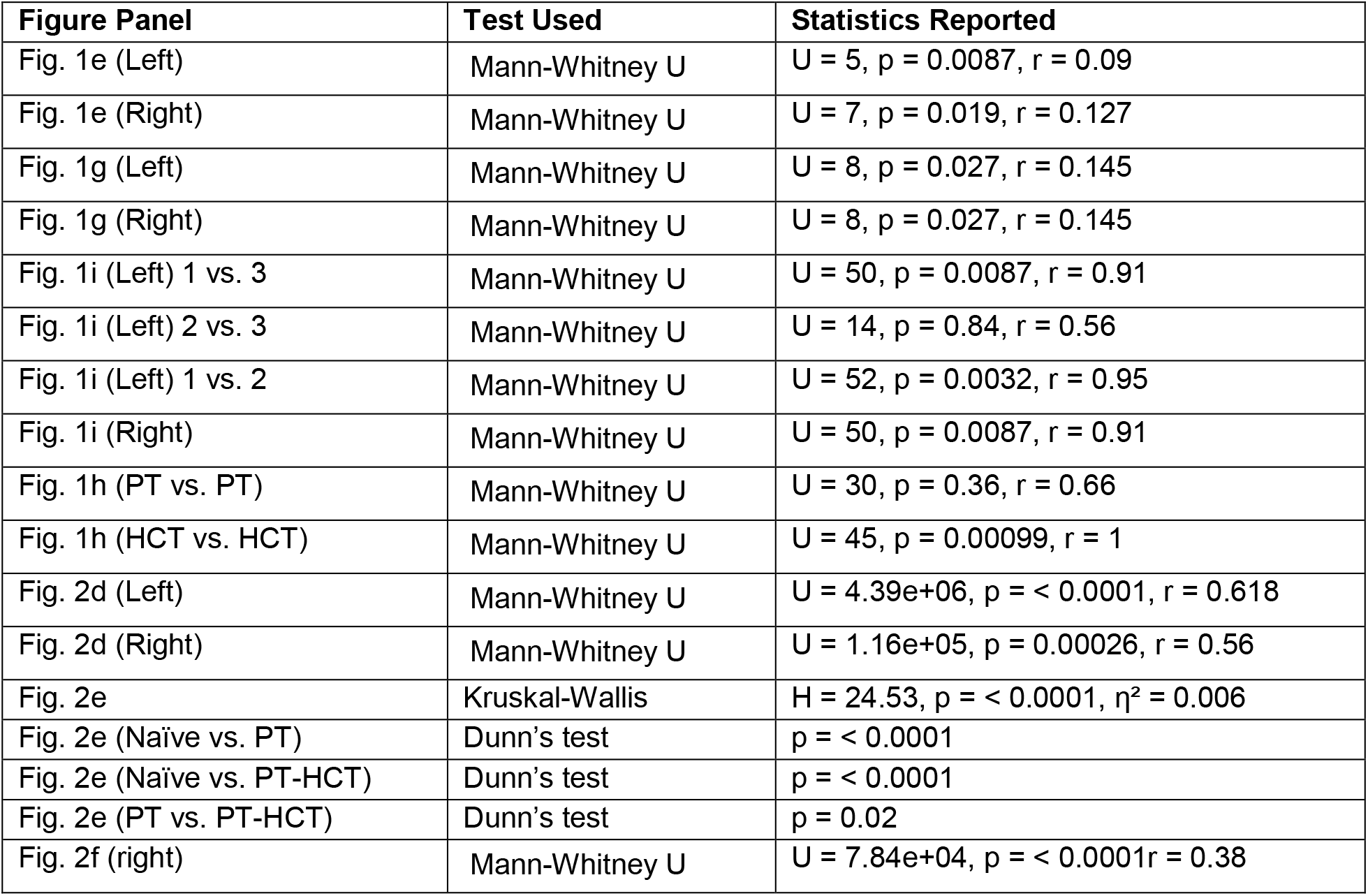
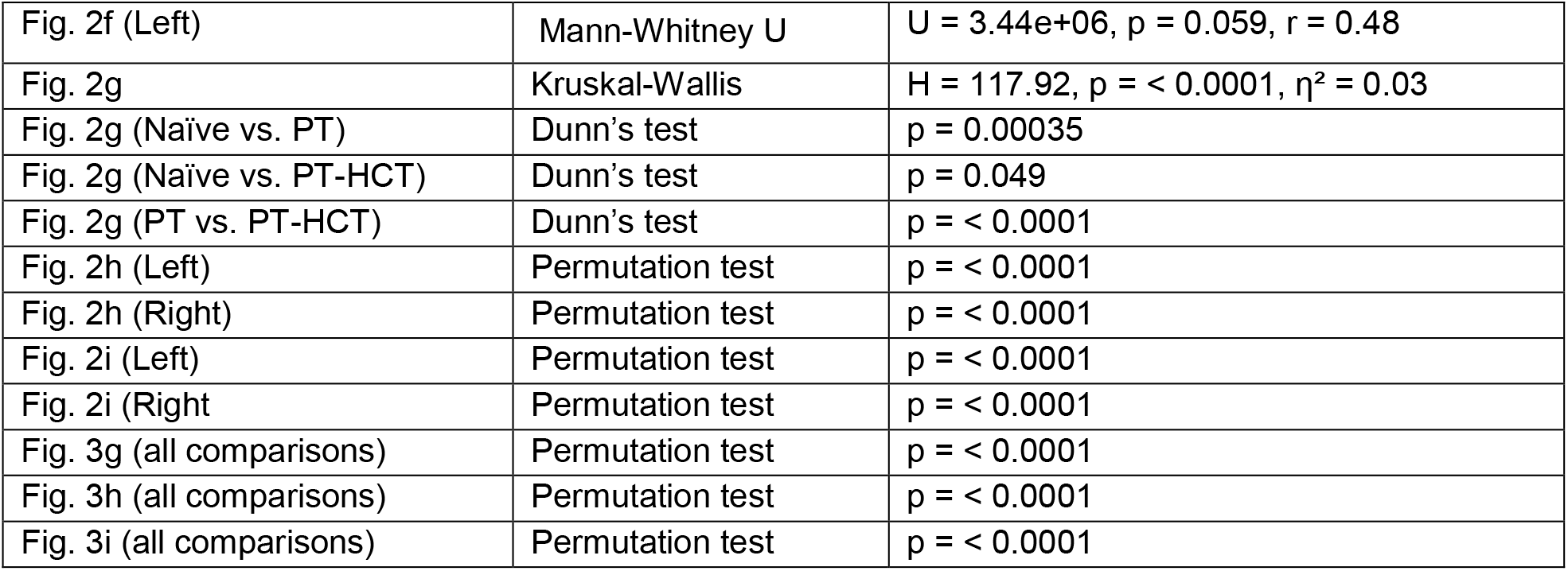
Summary of statistical tests and their outcomes for figures panels. The table includes test type, U-statistics, p-values, effect sizes (r), H-statistics and eta-squared (η^2^) where applicable.

| Figure Panel | Test Used | Statistics Reported |
| --- | --- | --- |
| Fig. 1e (Left) | Mann-Whitney U | $U = 5, p = 0.0087, r = 0.09$ |
| Fig. 1e (Right) | Mann-Whitney U | $U = 7, p = 0.019, r = 0.127$ |
| Fig. 1g (Left) | Mann-Whitney U | $U = 8, p = 0.027, r = 0.145$ |
| Fig. 1g (Right) | Mann-Whitney U | $U = 8, p = 0.027, r = 0.145$ |
| Fig. 1i (Left) 1 vs. 3 | Mann-Whitney U | $U = 50, p = 0.0087, r = 0.91$ |
| Fig. 1i (Left) 2 vs. 3 | Mann-Whitney U | $U = 14, p = 0.84, r = 0.56$ |
| Fig. 1i (Left) 1 vs. 2 | Mann-Whitney U | $U = 52, p = 0.0032, r = 0.95$ |
| Fig. 1i (Right) | Mann-Whitney U | $U = 50, p = 0.0087, r = 0.91$ |
| Fig. 1h (PT vs. PT) | Mann-Whitney U | $U = 30, p = 0.36, r = 0.66$ |
| Fig. 1h (HCT vs. HCT) | Mann-Whitney U | $U = 45, p = 0.00099, r = 1$ |
| Fig. 2d (Left) | Mann-Whitney U | $U = 4.39e+06, p = < 0.0001, r = 0.618$ |
| Fig. 2d (Right) | Mann-Whitney U | $U = 1.16e+05, p = 0.00026, r = 0.56$ |
| Fig. 2e | Kruskal-Wallis | $H = 24.53, p = < 0.0001, \eta^2 = 0.006$ |
| Fig. 2e (Naïve vs. PT) | Dunn's test | $p = < 0.0001$ |
| Fig. 2e (Naïve vs. PT-HCT) | Dunn's test | $p = < 0.0001$ |
| Fig. 2e (PT vs. PT-HCT) | Dunn's test | $p = 0.02$ |
| Fig. 2f (right) | Mann-Whitney U | $U = 7.84e+04, p = < 0.0001, r = 0.38$ |
| Fig. 2f (Left) | Mann-Whitney U | $U = 3.44e+06$, $p = 0.059$, $r = 0.48$ |
| Fig. 2g | Kruskal-Wallis | $H = 117.92$, $p = < 0.0001$, $\eta^2 = 0.03$ |
| Fig. 2g (Naïve vs. PT) | Dunn's test | $p = 0.00035$ |
| Fig. 2g (Naïve vs. PT-HCT) | Dunn's test | $p = 0.049$ |
| Fig. 2g (PT vs. PT-HCT) | Dunn's test | $p = < 0.0001$ |
| Fig. 2h (Left) | Permutation test | $p = < 0.0001$ |
| Fig. 2h (Right) | Permutation test | $p = < 0.0001$ |
| Fig. 2i (Left) | Permutation test | $p = < 0.0001$ |
| Fig. 2i (Right) | Permutation test | $p = < 0.0001$ |
| Fig. 3g (all comparisons) | Permutation test | $p = < 0.0001$ |
| Fig. 3h (all comparisons) | Permutation test | $p = < 0.0001$ |
| Fig. 3i (all comparisons) | Permutation test | $p = < 0.0001$ |

## Acknowledgments

We thank Pico Caroni, Mark Hübener and Flavio Donato for their valuable insights on the work. We thank Simon Saner from the electrical workshop for his support in building the behavior set-up. Funding: Research was supported by the Swiss National Science Foundation (310030-197850).

## Author contributions

Conceptualization, M.S., S.J., and T.R.B.; Methodology and experiments, S.J., M.N., and T.R.B.; Experimental support, M.N.; Analysis, M.S, S.J., T.R.B.; Original draft, M.S. and T.R.B.; Revisions, all authors; Supervision and funding, T.R.B.

